# *In vitro* assessment of the virucidal activity of four mouthwashes containing Cetylpyridinium Chloride, ethanol, zinc and a mix of enzyme and proteins against a human coronavirus

**DOI:** 10.1101/2020.10.28.359257

**Authors:** A. Green, G. Roberts, T. Tobery, C. Vincent, M. Barili, C. Jones

## Abstract

**Background:** saliva is established to contain high counts SARS-CoV-2 virus and contact with saliva droplets, contaminated surfaces or airborne particles are sources of viral transmission. The generation of infective aerosols during clinical procedures is of particular concern. Therefore, a fuller understanding of the potential of mouthwash to reduce viral counts and modulate the risk of transmission in medical professional and public context is an important research topic.

**Method:** we determined the virucidal activity of four anti-bacterial mouthwashes against a surrogate for SARS-CoV-2, Human CoV-SARS 229E, using a standard ASTM suspension test, with dilution and contact times applicable to recommended mouthwash use.

**Results:** the mouthwash formulated with 0.07% Cetylpyridinium Chloride exhibited virucidal effects providing a ≥3.0 log reduction HCoV-229E viral count. Mouthwashes containing 15.7% ethanol, 0.2% zinc sulphate heptahydrate and a mix of enzymes and proteins did not demonstrate substantive virucidal activity in this test.

**Conclusion:** mouthwash containing 0.07% Cetylpyridinium Chloride warrants further laboratory and clinical assessment to determine their potential benefit in reducing the risk of SARS-CoV-2.

**Highlights:** SARS-CoV-2 can be transmitted through contact with infective saliva.

Studies are needed to understand if mouthwash can lower SARS-CoV-2 transmission risk.

0.07% Cetylpyridinium Chloride (CPC) mouthwash exhibited virucidal effects against HCoV-SARS 229E.

Further studies on potential of 0.07% CPC mouthwash against SARS-CoV-2 are warranted.

## Introduction

One of the main transmission routes for SARS-CoV-2 is believed to be through aerosols and droplets of infective saliva, either directly via coughing, sneezing and exhalation or indirectly via fomites^1^.

SARS-CoV-2 saliva load has been reported to be up to 10^7^ copies per ml, suggesting active viral replication occurs in the mouth^2^. Salivary glands, oral mucosa and the tongue are reported to express the receptor angiotensin-converting enzyme, ACE-2, which SARS-CoV-2 uses to enter host cells^3, 4^. This suggests that reducing infective SARS-CoV-2 load in saliva could play a key role in reducing the risk of viral transmission. In addition to the general population, the risk of transmission from infective saliva aerosols generated during clinical procedures is of particular concern, including authors of a recent Cochrane review who urge researchers to conduct studies to understand the potential of anti-bacterial mouthwashes and sprays to protect healthcare professionals during aerosol-generating procedures^5, 6^. Therefore, an understanding of the potential of mouthwash use to reduce SARS-CoV-2 viral load and modulate the risk of transmission in professional and public settings is a pressing question to address.

Coronaviruses, including SARS-CoV-2, are enveloped viruses possessing a host derived lipid bilayer. This lipid bilayer can be disrupted by agents through lipid solubilisation, membrane disruption or damage to the embedded glycoproteins^7^. A recent publication reported that several commercial anti-bacterial mouthwashes possessed virucidal effects against SARS-CoV-2 *in vitro*, however, it is possible that other established anti-bacterial agents may have potential to inactivate enveloped viruses, including SARS-CoV-2^8^. In this paper we report the virucidal effect of four mouthwashes formulated with different anti-bacterial agents against a SARS-CoV-2 surrogate, Human Coronavirus Strain 229E, using a standard ‘time to kill’ suspension test and contact times appropriate to mouthwash use.

## Methods

The virucidal activity of each mouthwash was determined following the ASTM International Standard E1052-20 suspension protocol, *Standard Practice to Assess the Activity of Microbicides Against Viruses in Suspension*. Human Coronavirus Strain HCoV-229E (#VR-740) was sourced from ATCC (American Type Culture Collection) and high titer viral stocks were propagated and maintained by BioScience Laboratories Inc (Bozeman, Montana, USA). On the day of testing, stock virus was removed from a -70°C freezer and thawed prior to use with independent aliquots were prepared for each replicate. The host cell line, MRC-5 was obtained from ATCC (#CRL-171) and was maintained as monolayers in growth media containing 4% serum (FBS) at 35 +/- 2°C and 5% CO2 using disposable cell culture labware. 24 hours prior to testing, cells were seeded into multi-well cell culture plates. For viral infectivity testing, cell monolayers were used at 80-90% confluence and were maintained under serum-free conditions. Prior to testing, neutralisation studies of each product were conducted versus the test virus to ensure the virucidal properties of the test product were effectively neutralised and the neutraliser solution was non-toxic to both the challenge virus and the host cell line. For all test mouthwash products, Dey-Engley neutralising broth (D/E broth) was effective and used in all testing. Test solutions were further detoxified by passing them through a Sephadex column.

Neutralisation and Cytotoxicity controls were performed in tandem with testing and were used as controls in analysis of the results.

For the test treatment, a 0.5ml of viral aliquot was added to 4.5ml of undiluted mouthwash and incubated for 30 and 60 seconds at ambient temperature. All test treatments were conducted in triplicate and plating was performed with four replicates. The post exposure infectivity TCID50 (50% tissue culture infectious dose) was determined using the Quantal test (Spearman -Kärber method) and a mean log10 reduction was calculated as the difference in TCID50. Test mouthwashes were:

Mouthwash containing 0.07% Cetylpyridinium Chloride (CPC), sodium fluoride, and flavour oil

Mouthwash containing 15.7% ethanol, sodium fluoride, and flavour oil.

Mouthwash containing 0.2% zinc sulphate heptahydrate, sodium fluoride and flavour oil.

Mouthwash containing mix of Amyloglucosidase, Glucose Oxidase, Lysozyme, Colostrum, Lactoferrin, Lactoperoxidase, sodium fluoride and flavour oil.

## Results

Mouthwashes are water based formulations, containing humectants, emulsifiers, a fluoride source, preservatives, and flavour oils to provide pleasant taste. To this similar formulation chassis, anti-bacterial agents are incorporated. In this study, we compared four anti-bacterial mouthwashes containing actives hypothesised to act against enveloped viruses. CPC is a quaternary ammonium compound which interacts with the viral envelope through cationic charge-based interaction leading to gross distortion of the viral ultrastructure^9^. Similarly, alcohols including ethanol, perturb the lipid membrane leading to leakage^7^. Zinc cations are reported to inhibit replication of SARS-CoV and binding of SARS-CoV-2 to AC2-E receptors^10, 11^. The mouthwash with enzymes and proteins is designed to generate hypothiocyanate from the Lactoperoxidase (LPO) system, with the LPO system reported to possess virucidal activity against a number of enveloped viruses^12^. Additionally, lysozyme and lactoferrin are components of innate viral defences, and as cationic proteins, can be hypothesised to act on the viral lipid envelope^13^.

However, of these four anti-bacterial mouthwashes, only the mouthwash containing 0.07% CPC exhibited viricidal activity against HCoV-229E, providing a mean 3.08 TCID50 log10 reduction in viral count. This corresponds to ≥99.9% reduction in viral count and was achieved with contact times relevant to mouthwash use. Contact with ethanol, zinc and enzyme and protein mouthwashes did not provide substantial reduction in viral counts in this *in vitro* test. A summary of results for all mouthwashes is provided in Table 1.

**Table 1.** Virucidal efficacy of mouthwashes against HCoV-229E.

| **Mouthwash** | **Contact Time (seconds)** | **Replicate** | **TCID<sub>50</sub> Reduction-log10** | **Mean TCID<sub>50</sub> Reduction-log10 (SD)** |
| --- | --- | --- | --- | --- |
| CPC | 30 | 1 | $\geq 3.25$ | 3.08(0.29) |
| | | 2 | $\geq 3.25$ | |
| | | 3 | $\geq 2.75$ | |
| | 60 | 1 | $\geq 3.25$ | 3.08(0.29) |
| | | 2 | $\geq 3.25$ | |
| | | 3 | $\geq 2.75$ | |
| Ethanol | 30 | 1 | 0.50 | 0.17(0.29) |
|  |  | 2 | 0.00 |  |
|  |  | 3 | 0.00 |  |
|  | 60 | 1 | 0.50 | 0.33(0.29) |
|  |  | 2 | 0.50 |  |
|  |  | 3 | 0.00 |  |
| Zinc | 30 | 1 | 1.25 | 1.17(0.38) |
|  |  | 2 | 1.50 |  |
|  |  | 3 | 0.75 |  |
|  | 60 | 1 | 2.00 | 1.83(0.14) |
|  |  | 2 | 1.75 |  |
|  |  | 3 | 1.75 |  |
| Enzymes and proteins | 30 | 1 | 0.25 | 0.25(0.25) |
|  |  | 2 | 0.50 |  |
|  |  | 3 | 0.00 |  |
|  | 60 | 1 | 0.25 | 0.25(0.25) |
|  |  | 2 | 0.50 |  |
|  |  | 3 | 0.00 |  |

## Discussion

Currently, there are few published studies reporting efficacy of anti-bacterial mouthwashes against SARS-CoV-2 or surrogates. This is against a backdrop of the COVID-19 pandemic with medical professionals urging research into the potential benefits of mouthwash to reducing viral load and lower risk of transmission^5^. In this study, we confirm that not all anti-microbial mouthwashes possess virucidal benefits. Against a SARS COV2 surrogate HCoV-229E, at relevant in use contact times, only the 0.07% CPC mouthwash proved to be effective in reducing viral count by greater than 99.9%. Though preliminary, this result agrees with evidence of CPC activity against influenza virus and a recently published clinical trial with COVID-19 patients, where a 0.075% CPC containing mouthwash was reported to reduce saliva viral load for up to 6 hours after use^9,14^. Contact with the mouthwash containing 15.7% ethanol did not lead to significant reduction in count, and given alcohol levels of 60% - 95% v/v are used in hand hygiene products, it seems likely that a mouthwash will require a higher concentration of ethanol or combination of agents to bring about a reduction in viral load^15^.

Under the conditions of this test, mouthwashes with zinc and the enzyme and proteins mix did not substantively impact viral count, however, these agents are reported to have viricidal action and merit further study. In summary, based on our findings, we propose the need further *in vitro* and clinical assessment of potential of 0.07% CPC mouthwash in the management of SARS-CoV-2 transmission in clinical and public settings.

## Acknowledgements

The laboratory study was conducted by BioScience Laboratories Inc. We are grateful to Andrew Joiner, Sayandip Mukherjee and Tamsin Worrad-Andrews for useful discussions.

## Ethics

Not applicable to this *in vitro* study.

## Author contributions

Glyn Roberts conceived the study, provided interpretation of data, revised the draft, and approved the final version. Alison Green provided interpretation of data, drafted, and wrote the paper, and approved the final version. Carol Vincent designed the work, contributed to the acquisition and analysis, interpretation of data, revised the draft, and approved the final version. Timothy Tobery designed the work, contributed to the acquisition and analysis, and interpretation of data, revised the draft, and approved the final version. Matteo Barili contributed to the acquisition of data, revised the draft, and approved the final version. Carolyn Jones contributed to the interpretation of data, revised the draft, and approved the final version.

